# BDNF Regulates Pituitary Stem Cell Engagement towards precursor state

**DOI:** 10.64898/2026.04.02.716194

**Authors:** Kevin Sochodolsky, Konstantin Khetchoumian, Aurelio Balsalobre, Ryan M. Feeley, Margaret E. Rice, Probir Chakravarty, Robin Lovell-Badge, Karine Rizzoti, Jacques Drouin

## Abstract

Following their engagement towards differentiation, tissue stem cells often transit through a precursor state that is difficult to define because of its transient nature; similarly, the precise role of lineage precursors in implementation of tissue architecture and function is unknown. In the present work, we used two mouse models of deficient feedback regulation to characterize precursors of the pituitary corticotrope lineage that regulates the stress response. Both the POMC knockout and adrenalectomized mouse models develop glucocorticoid deficiency and compensatory accumulation of corticotrope precursors that have so far eluded characterization. We found that pre-corticotrope differentiation depends on the lineage-specific factor Tpit and is repressed by glucocorticoids. We identified brain-derived neurotrophic factor (BDNF) as the signal that engages pituitary stem cells towards differentiation in these models as well as in normal pituitary development. A glucocorticoid-sensitive BDNF autocrine loop active in pre-corticotropes turns these cells into signaling hubs for maintenance of pituitary-adrenal homeostasis.

**Highlights:**

- Pituitary lineage precursors expand in conditions of deficient feedback regulation
- BDNF mobilizes pituitary stem cells during establishment of tissue size and architecture
- Corticotrope precursors are a signaling hub for tissue homeostasis

## Introduction

During development, many multicellular organs derive from a population of stem cells that give rise to different lineages. Cells of these different lineages then organize into stereotypic arrangements that define the organ’s cellular architecture. The pituitary gland has been an interesting model in this respect in that it does not have a striking cellular organization and superficially appears to be a random patchwork of cells. This simple view was contradicted when specific homotypic and heterotypic cell-cell interactions were identified and found to adapt to various physiological conditions (Mollard, Hodson et al. 2012). Thus, dynamic tissue modeling in development (Bonnefont, Lacampagne et al. 2005, Budry, Lafont et al. 2011) and remodeling under various physiological conditions (Lafont, Desarmenien et al. 2010, Sanchez-Cardenas, Fontanaud et al. 2010, Hodson, Schaeffer et al. 2012) occur to adapt pituitary architecture. During these processes, new cells arise through differentiation from a pool of tissue stem cells. Pituitary stem cells (PSCs) were identified in the developing gland and an adult pool of PSCs is retained along the margin of the cleft that separates the anterior from intermediate pituitary lobes (Fauquier, Rizzoti et al. 2008, Drouin and Brière 2022). PSCs are also present in the parenchyma of the anterior lobe. Whether during development or in remodelling conditions in the adult, the signals that orchestrate pituitary homeostasis for maintenance of the proportion of the different lineages remain largely unknown.

While the identification of transcription factors involved in cell specific lineage establishment helped to define transcriptional regulatory mechanisms that direct terminal differentiation of different pituitary lineages (Drouin and Brière 2022), the mechanisms that trigger PSC differentiation towards specific lineages are poorly understood. PSCs are marked by the expression of SOX2, SOX9 and additional markers according to their localization in the gland. (Chen, Gremeaux et al. 2009, Rizzoti, Chakravarty et al. 2023). Proliferation of PSCs progeny in young animals involves paracrine Wnt signalling (Russell, Lim et al. 2021), but signals for engagement of PSCs towards differentiation are still missing. It has long been known that hormones originating from target tissues exert feedback on pituitary cells including on their number. Thus, end-organ ablation triggers an expansion of related pituitary lineages such an increase in ACTH-producing corticotrope cell number in response to adrenalectomy (ADX) (Nolan and Levy 2006). In this instance, mobilization of PSCs was shown in contrast to basic tissue maintenance of corticotropes that depends primarily on self-duplication of differentiated corticotropes (Langlais, Couture et al. 2013). The early studies suggested that a common or similar pool of precursors originating from PSCs may be involved in lineage expansion in response to ADX (Nolan and Levy 2006). More detailed analyses using the ADX model first confirmed this (Rizzoti, Akiyama et al. 2013), and subsequently showed that while mobilization and expansion of PSCs is initially non-cell type specific, progeny of the appropriate lineage is selectively maintained (Rizzoti, Chakravarty et al. 2023). In the present work, we addressed underlying mechanisms using another model of end-organ deficit, the *POMC* knockout (*POMC^-/-^*) mice, that present with a large hyperplasia of cells of the corticotrope lineage (Yaswen, Diehl et al. 1999). This led us to define in both models similar nascent corticotrope precursors, the pre-corticotropes, and to identify BDNF as a signalling pathway for mobilization of PSCs and maintenance of the pre- corticotrope fate. During pituitary gland development, the BDNF pathway is not important for fetal establishment of the gland but rather, it is critical for postnatal expansion of the pituitary that occurs in the first postnatal weeks in mice (Laporte, Vennekens et al. 2020), leading to a smaller pituitary in *BDNF* knockout (*BDNF^-/-^*) mice. As all lineages appear to be affected by BDNF deletion, this signalling appears primarily responsible for mobilization of PSCs during tissue expansion.

## Results

### *POMC* knockout mice develop glucocorticoid and Tpit dependent pituitary hyperplasia

Consistent with end-organ ablation, adrenalectomy (ADX) in this case (Nolan and Levy 2006), mice inactivated for the *POMC* gene exhibit marked pituitary hyperplasia of POMC-expressing corticotrope cells late in life (Yaswen, Diehl et al. 1999). These mice have adrenal hypoplasia resulting from the lack of POMC-derived ACTH (Coll, Challis et al. 2004). In this respect, they might be viewed as similar to ADX mice, but the reported pituitary hyperplasia appeared far more severe. We reinvestigated this model to assess the mechanisms for engagement of PSCs in this context. As reported, old (fully penetrant in both sexes) *POMC^-/-^*mice develop extensive hyperplasia: most hyperplastic cells are positive for the POMC lineage marker Tpit (Figure 1A, B) that normally drives terminal differentiation of the POMC lineages (Lamolet, Pulichino et al. 2001, Pulichino, Vallette-Kasic et al. 2003, Pulichino, Vallette-Kasic et al. 2003). As the adrenals fail to develop in absence of ACTH, the *POMC^-/-^* mice are hypocortisolemic (Yaswen, Diehl et al. 1999) and the absence of glucocorticoids (Gc) may play a role in pituitary hyperplasia. To test this, we provided the synthetic Gc dexamethasone (Dex) in the drinking water of *POMC^-/-^*mice for 2 weeks: this resulted in significant reduction of hyperplastic cell number as assessed by Tpit labelling (Figure 1C, D). The hyperplastic pituitary also exhibits a significant increase in Ki67-positive dividing cells (Figure 1B, E) and this is completely abrogated by Dex administration (Figure 1E). The suppressive effect of Dex may result from both its negative feedback action and apoptosis of Ki67-positive cells (Nolan and Levy 2006).

**Figure 1.**
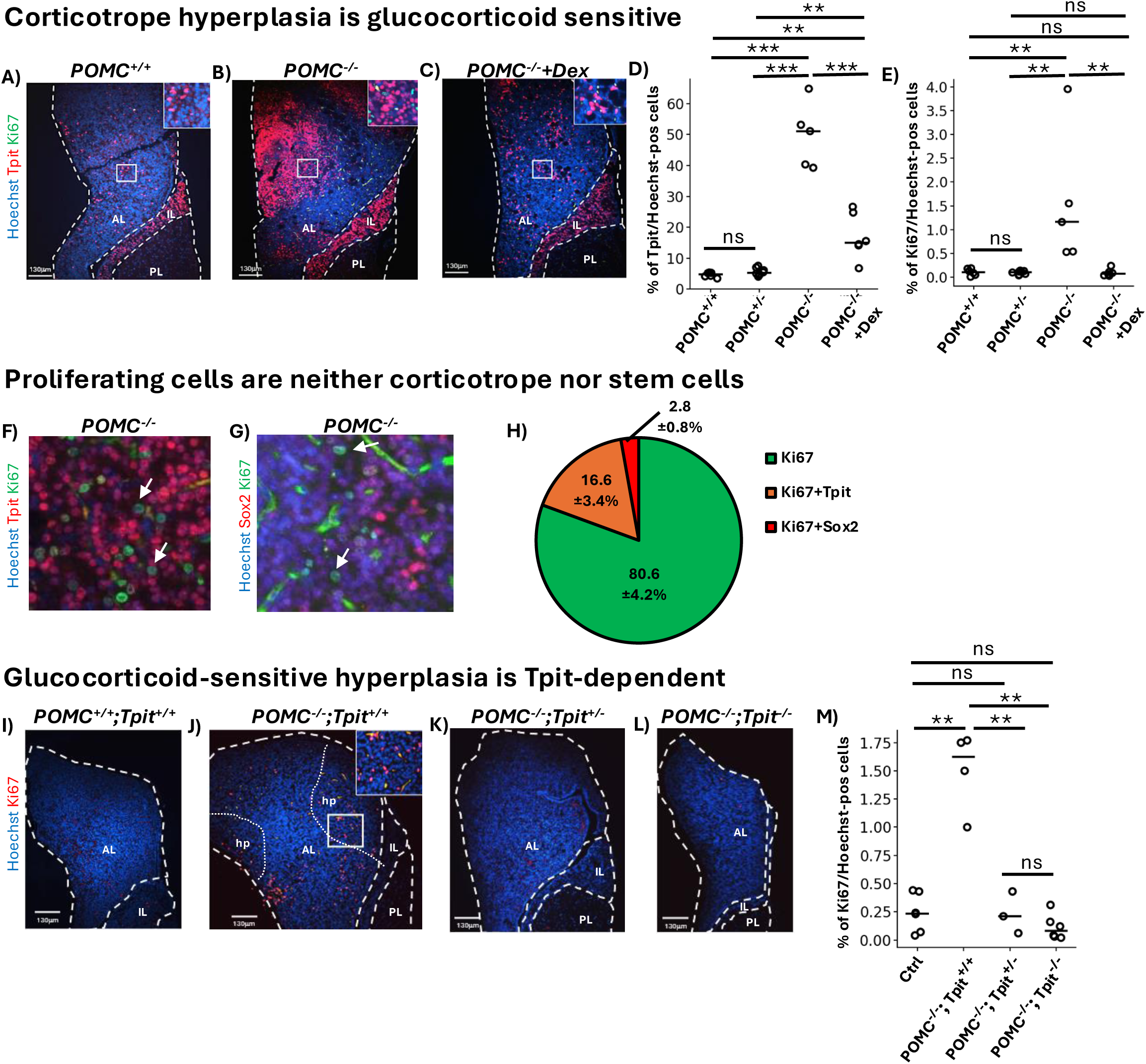
*POMC* knockout mice develop glucocorticoid and Tpit sensitive pituitary hyperplasia. (**A-E**) Corticotrope hyperplasia is glucocorticoid sensitive. Photomicrographs of mouse pituitary sections stained for Tpit and Ki67 reveal Tpit-positive hyperplasia in *POMC* knockout pituitary (**B**, *POMC^-/-^)* compared to normal pituitary (**A**, *POMC^+/+^*). Administration of dexamethasone (Dex) to *POMC^-/-^* mice decreases Tpit-positive hyperplasia (**C**). The reversal of hyperplasia by Dex treatment was quantitated as the number of Tpit positive cells (*POMC^+/+^* n=6 from 6 mice, *POMC^+/-^* n=8 from 8 mice, *POMC^-/-^* n=5 from 5 mice & *POMC^-/-^*+Dex n=6 from 6 mice) (**D**) and the number of Ki67 positive cells (*POMC^+/+^* n=6 from 6 mice, *POMC^+/-^*n=8 from 8 mice, *POMC^-/-^* n=5 from 5 mice & *POMC^-/-^*+Dex n=6 from 6 mice) (**E**) (**F-H**) Ki67 positive proliferating cells are neither corticotropes as revealed by co-staining for Ki67 and Tpit (**F**), nor pituitary stem cells as revealed by co-staining for Sox2 and Ki67 (**G**). (**H**) Quantitation of single or double staining for either Tpit, Sox2 and/or Ki67 as indicated (n=4 from 4 mice). Error corresponds to SEM. (**I-M**) Dex sensitive hyperplasia is Tpit-dependent. The hyperplasia (hp) and Ki67-positive cells observed in *POMC^-/-^* ;*Tpit^+/+^* pituitary (**J**) compared to wildtype (*POMC^+/+^*;*Tpit^+/+^*) pituitary (**I**) is reversed in Tpit heterozygote (*POMC^-/-^*;*Tpit^+/-^*) (**K**) and homozygote (*POMC^-/-^*;*Tpit^-/-^*) (**L**) knockout mice. (**M**) Quantitation of the number of Ki67 positive cells in pituitaries of the indicated genotypes (Ctrl n=6 from 6 mice, *POMC^-/-^;Tpit^+/+^* n=4 from 4 mice, *POMC^-/-^;Tpit^+/-^*n=3 from 3 mice & *POMC^-/-^;Tpit^-/-^* n=6 from 6 mice). Statistical significance was measured by paired two-sided T-test: * p≤0.05, **p≤0.01, ***p≤0.001.

It is noteworthy that Ki67-positive cells of *POMC^-/-^*mice are mostly negative for Tpit as assessed by immunofluorescence (IHF) colabeling (Figure 1F, H). We speculated that they may be dividing PSCs and performed co-labelling for SOX2 and Ki67: the Ki67-positive cells are rarely positive for SOX2 (Figure 1G, H) and may therefore represent precursor cells transiting between PSCs and differentiation.

Interestingly, *Tpit*^-/-^ mice fail to differentiate and expand the POMC lineages leading to adrenal hypoplasia and hypocortisolism like *POMC^-/-^* mice (Pulichino, Vallette-Kasic et al. 2003) but without pituitary hyperplasia. Having observed that the pituitary hyperplasia present in *POMC^-/-^*mice is Dex-dependent (Figure 1C-E), we wondered why the *Tpit*^-/-^mice do not develop pituitary hyperplasia. We therefore crossed the *Tpit* and *POMC* knockout mice and found that the *POMC^-/-^* hyperplasia (Figure 1I, J) is completely prevented by inactivation of the *Tpit* gene, whether one (Figure 1K) or two (Figure 1L) copy of the gene is inactivated. Prior work has identified Tpit target genes that are either dosage dependent or not (Khetchoumian, Sochodolsky et al. 2024). Since Tpit is essential for corticotrope differentiation, it is not surprising that corticotrope cells are not detected in the double knockout mice. However, these mice could have developed a hyperplastic response of undifferentiated cells, but labelling with Ki67 indicated that this is not the case (Figure 1M). Since *Tpit* heterozygosity does not prevent corticotrope differentiation (Pulichino, Vallette-Kasic et al. 2003), the *POMC^-/-^;Tpit^+/-^* mice could have expanded this lineage but this was not observed (Figure S1). In summary, *POMC^-/-^* mice develop pituitary hyperplasia of cells that are positive for the corticotrope marker Tpit and that appear to result from expansion of a population of transient cells that may represent a pre-corticotrope fate.

### Single cell transcriptomics identifies a pre-corticotrope cell identity

To define the nature of the hyperplastic and Ki67-positive dividing cells, we performed single-cell transcriptome (scRNA-seq) analyses with *POMC^-/-^* compared to *POMC^+/+^* pituitaries (Figure 2A, B). The *POMC^-/-^* pituitary is greatly enriched in Tpit-positive cells (Figure 2B arrow). These corticotrope-like cells subdivide into two major clusters that now account for about 65% of the cells compared to 10% for the *POMC^+/+^* pituitary. We used pseudotime analysis (Cao, Spielmann et al. 2019) of the corticotrope-related and PSC lineages to define their relationship: starting with PSCs, the lineage goes through Ki67-positive proliferating cells towards presumptive pre-corticotropes and corticotropes (Figure 2C). Consistent with the IHF analyses (Figure 1F-H), the Ki67-positive cells (Figure 2D) are negative for Sox2 (Figure 2E) and Sox9 (Figure 2F) whereas both pre-and corticotropes are positive for Tpit (Figure 2G) consistent with the interpretation that Ki67-positive cells represent proliferating transit cells. Differential gene expression analyses of these different cell clusters identified keratins 8 and 18 for their expression in PSCs (Cox, Laporte et al. 2019) and pre-corticotropes but not in corticotropes (Figure 2H). Identification of the corticotrope and pre-corticotrope sub-clusters is further supported by the expression patterns of Tpit and CRH receptor (CRHR1) (Figure S2). Whereas PSCs of *POMC^+/+^*and *POMC^-/-^* animals have similar gene expression signatures, the proliferating cells of the *POMC^-/-^* pituitary retain the Krt8 and Krt18 markers of PSCs while showing weak gains of corticotrope markers (Figure 2I). Krt18 expression was ascertained in the *POMC^-/-^* pituitary by IHF and it is clearly associated with the hyperplastic cells (Figure 2J). These data suggest that *POMC^-/-^*PSCs are mobilised into proliferation and differentiation and accumulate as large pre-corticotrope and corticotrope cell clusters.

**Figure 2.**
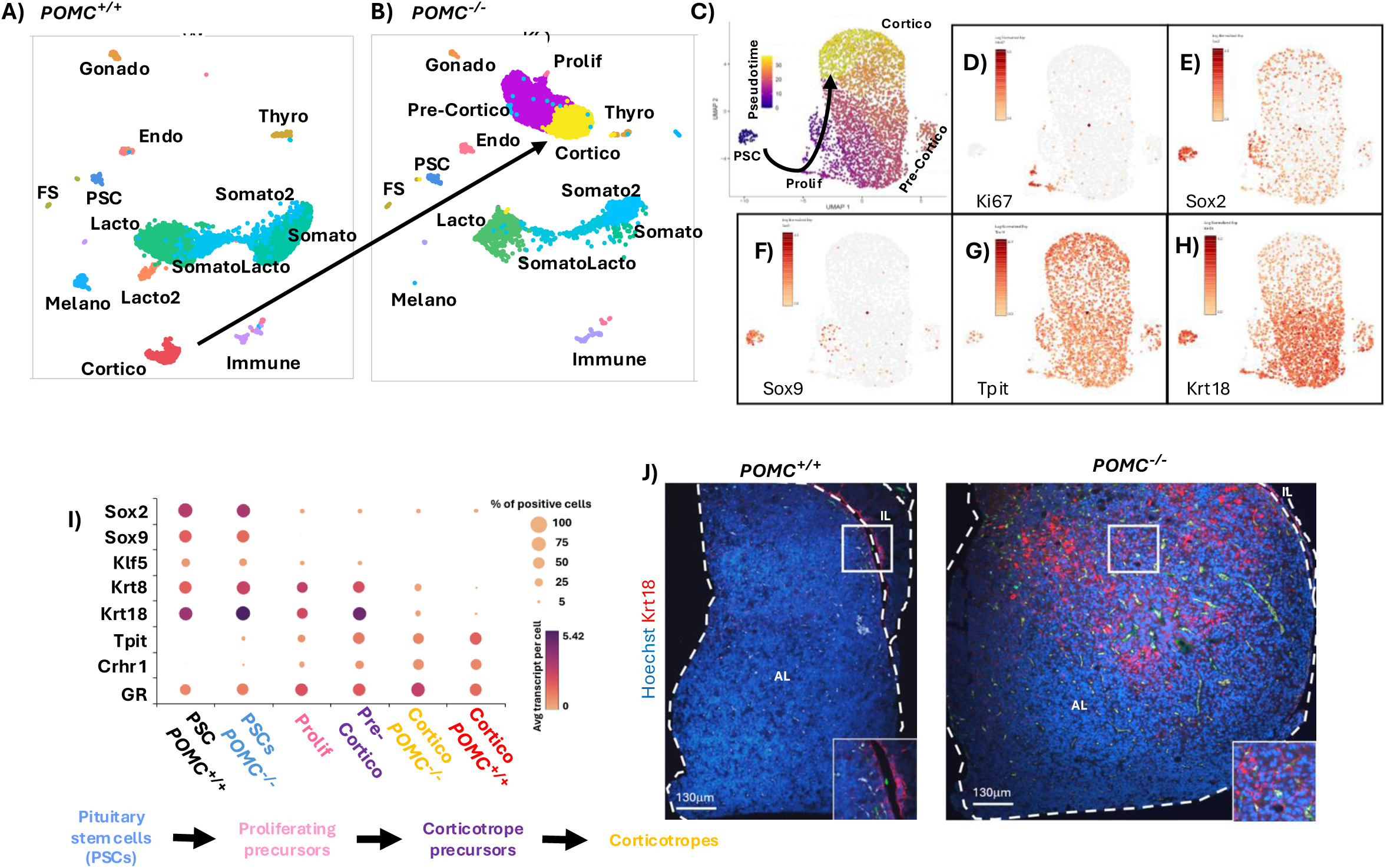
Single cell RNA-Seq (scRNA-seq) analyses of pituitary cells reveal subset of pre-corticotrope cells. (**A, B**) Aggregate analysis of scRNA-seq data from wildtype (*POMC^+/+^*) and knockout (*POMC^-/-^*) pituitaries reveal shift of *POMC^+/+^*corticotrope cluster to a new cluster in *POMC^-/-^* pituitaries (**B**) as indicated by arrow. (**C**) Pseudotime analysis of the isolated pituitary stem cell (PSC) and corticotrope clusters of *POMC^-/-^*mice indicate progression from PSCs through Ki67-positive proliferating cells towards pre-corticotropes and corticotropes. (**D-H**) Identification of sub-clusters within the *POMC^-/-^* corticotrope cluster identifies proliferating cells by Ki67 expression (**D**) whereas Sox2 and Sox9 expression (**E-F**) highlight PSCs, and Tpit expression labels the pre-corticotrope and corticotrope sub-clusters (**G**). The pre-corticotrope sub-cluster is further highlighted by expression of keratin18 (Krt18, **H**). (**I**) Dotplot visualization of relative expression for the indicated genes in the various sub-clusters as indicated. The color of the dot indicates relative expression levels, and the size is the proportion of cells expressing said gene. (**J**) Immunohistofluorescence (IHF) for Krt18 on sections of pituitaries from *POMC^+/+^* and *POMC^-/-^* mice.

### BDNF, a candidate trophic factor for pre-corticotrope expansion

To identify candidate trophic factors that may be produced by pre-corticotropes to engage PSCs into proliferation and differentiation, we analyzed differentially expressed genes between pre-corticotropes and corticotropes (Figure 3A, Table S1): BDNF stood out as the most differentially expressed ligand-coding gene. Further support for this candidate came from CellChat analyses that identified the BDNF-*Ntrk2* ligand-receptor pair for interactions between pre-corticotropes and PSCs (Figure 3B). Indeed, BDNF is most expressed in pre-corticotropes compared to other cells of either *POMC^+/+^* or *POMC^-/-^*pituitaries (Figure 3C, Figure S3A). The BDNF receptor, TrkB *(Ntrk2)*, is expressed in many cells including PSCs, pre-corticotropes and corticotropes (Figure 3D, Figure S3B). Finally, the BDNF-responsive gene *Vgf* (Salton, Ferri et al. 2000) is most active in pre-corticotropes and proliferating cells of *POMC^-/-^* pituitaries with lower expression in corticotropes and PSCs (Figure 3E, Figure S3C) consistent with responsiveness of these cells to BDNF. With regards to PSCs, we found few double Sox2- and Ki67-positive cells suggesting that their entry into cycle may be associated with extinction of Sox2 expression; the *POMC^-/-^* PSCs show higher expression of cyclin D1 consistent with this model (Figure 3F). In summary, the BDNF-TrkB ligand-receptor pair is the strongest one identified by CellChat that is unique to the *POMC^-/-^* pre-corticotrope-PSC pair (Figure 3B).

**Figure 3.**
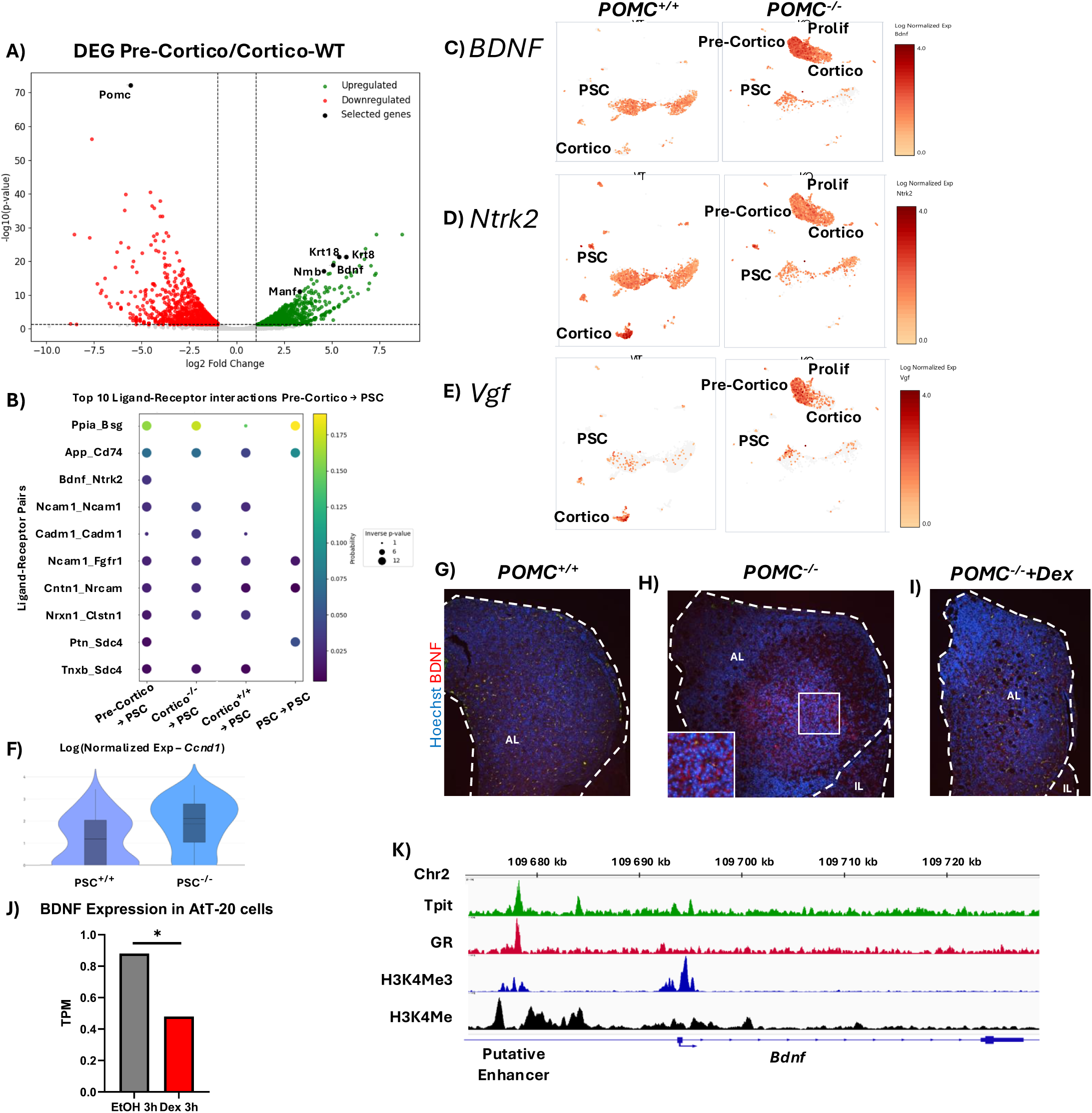
Dexamethasone-sensitive paracrine signalling between pre-corticotropes and pituitary stem cells. (**A**) Volcano plot showing differentially expressed genes (DEG) in pre-corticotropes (*POMC^-/-^*) compared to corticotropes (*POMC^+/+^*). The complete list of DEGs is provided in Supplementary Table 1. (**B**) CellChat analysis of candidate ligand-receptor pairs active between various pituitary cells as revealed by analysis of scRNA-seq data. Dot color represents probability score, and dot size inversely correlates with p-value. (**C-E**) UMAP representation of *BDNF* (**C**), *Ntrk2* (**D**) and *Vgf* (**E**) expression in *POMC^-/-^* and *POMC^+/+^*scRNA-seq datasets. (**F**) Violin plot of relative Cyclin D1 expression in *POMC^+/+^* and *POMC^-/-^* PSCs. (**G-I**) Immunofluorescence (IHF) staining of BDNF expression in pituitary sections from *POMC^+/+^* (**G**), *POMC^-/-^* (**H**) and *POMC^-/-^*mice treated for 2 weeks by Dex (**I**). (**J**) Glucocorticoid repression of *BDNF* gene expression in AtT-20 cells treated for 3 hours with Dex is revealed in RNA-Seq data from (Langlais, Couture et al. 2012), *p≤0.05. (**K**) Gene browser view of ChIP-seq data for the indicated marks around the *BDNF* locus. The ChIP-seq data are from previously published work (Budry, Balsalobre et al. 2012, Langlais, Couture et al. 2012, Mayran, Khetchoumian et al. 2018).

Prior work had shown Gc regulation of the *BDNF* gene in neurons (Chen, Lombès et al. 2017) and hence we interrogated the effect of Dex on BDNF expression in *POMC^-/-^*pituitaries by IHF. BDNF is expressed in the hyperplastic mass of *POMC^-/-^* compared to *POMC^+/+^*pituitaries (Figure 3G and H) and this expression is blocked by Dex treatment of *POMC^-/-^*mice (Figure 3I). To investigate the molecular basis of BDNF action on POMC expression, we queried our RNA-Seq data from the corticotrope model AtT-20 cells (Langlais, Couture et al. 2012): Dex represses BDNF expression in these cells (Figure 3J). We used our extensive ChIP-seq data in AtT-20 cells (Budry, Balsalobre et al. 2012, Langlais, Couture et al. 2012, Mayran, Khetchoumian et al. 2018) to identify a GR recruitment site located ∼15 Kb upstream of the *BDNF* gene that also recruits the corticotrope-restricted transcription factor Tpit; this site has the hallmark H3K4me1 signature of an active enhancer (Figure 3J). Prototypic DNA binding sites for GR (Langlais, Couture et al. 2012) and Tpit (Budry, Balsalobre et al. 2012) are present under their ChIP-seq peaks (not shown).

Collectively, these data suggests that BDNF produced by pre-corticotropes may act as a trophic factor to engage PSCs into proliferation and differentiation towards the pre-corticotrope fate, and that it may provide an autocrine loop for maintenance of pre-corticotropes.

### Adrenalectomy induces BDNF signalling from pre-corticotropes

We previously used a PSC lineage tracing model to show that ADX induces recruitment of PSCs for expansion of corticotrope cells (Rizzoti, Akiyama et al. 2013, Rizzoti, Chakravarty et al. 2023): we thus asked whether pre-corticotropes are expanded following ADX and whether they exhibit a similar transcriptional signature including BDNF expression as in the *POMC^-/-^* model. For this purpose, we analyzed the cell responses 48 hours after ADX in a scRNA-seq dataset (Rizzoti, Chakravarty et al. 2023) where we had integrated whole glands and PSC-enriched Sox9iresGFP positive fractions from both sham-operated and ADX males (Figure4A). Following ADX, a clearly discernable cluster of pre-corticotropes is present in scRNA-seq analyses (Figure 4B). We previously showed that GFP persistence from the Sox9iresGFP allele allows for a short-term tracing of PSCs (Rizzoti, Chakravarty et al. 2023). Here we show that pre-corticotropes induced after ADX are comprised in the Sox9iresGFP positive dataset, strongly suggesting that these cells originate from PSCs (Figure 4A and B). To assess putative ligand:receptor interactions in this model, we isolated the corticotrope and PSC clusters and performed CellChat analyses (Figure 4C). These analyses suggested particularly strong interactions involving BDNF and *Ntrk2* receptor between pre-corticotropes and the anterior lobe parenchymal SC-ALDH1A2 subset (Rizzoti, Chakravarty et al. 2023) of PSCs. Similar signalling is also suggested for pre-corticotropes and other PSC subtypes. UMAP projections of *Bdnf*, *Ntrk2* and *Vgf* expression in the scRNA-seq analyses of ADX compared to sham control mice indicate that *Bdnf* is most expressed in ADX-dependent pre-corticotropes whereas the *Ntrk2* receptor is expressed in most cells (Figure 4E, Figure S4 A and B). Interestingly, the BDNF-induced *Vgf* gene that indicates activity of the BDNF pathway is most expressed in pre-corticotropes (Figure 4F, Figure 4C). This interpretation is supported by RNA-scope analyses of *Vgf* expression on pituitary sections: *Vgf* colocalizes with POMC exclusively after ADX (Figure 4G) consistent with expression in nascent pre-corticotropes. A specific PSC subcluster exhibits increased cyclin D1 expression following ADX (Figure 4D) consistent with the *POMC^-/-^* model (Figure 3F).

**Figure 4.**
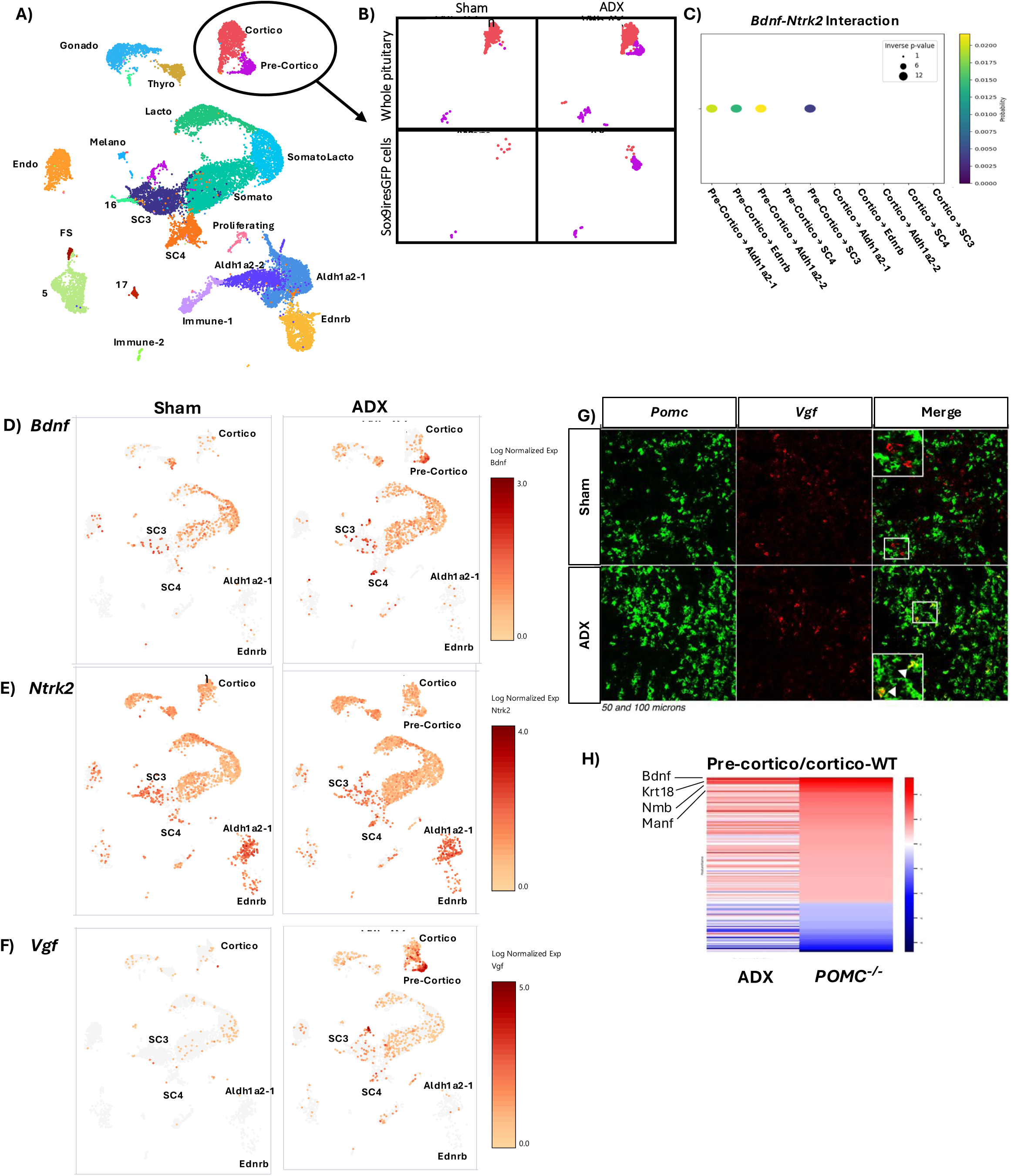
Adrenalectomy (ADX) induces BDNF signalling from pre-corticotropes. (**A**) UMAP representation of aggregate single cell RNA-Seq data for whole pituitaries combined with purified Sox9iresGFP pituitary progenitors. (**B**) Isolated corticotrope and Sox9+ PSC clusters from ADX and Sham operated mice revealing an ADX-dependent cluster of pre-corticotropes (purple). (**C**) CellChat analysis of BDNF-Ntrk2 ligand-receptor pair active between different corticotrope and PSC clusters. Dot color represents probability score, and dot size inversely correlates with the p-value. (**D-F**) UMAP representations of the corticotrope and PSC subclusters highlighting expression of the *BDNF* (**D**) *Ntrk2* (**E**) and *Vgf* (**F**) genes. (**G**) RNAscope analysis of *Vgf* gene expression in normal (Sham) and ADX pituitaries showing colocalization of VGF and POMC mRNA in ADX pituitaries. (**H**) Heatmap comparison of genes differentially expressed in pre-corticotropes compared to mature corticotropes identified in the ADX model compared with the *POMC^-/-^*model.

Direct comparison of gene expression profiles for pre-corticotropes defined in the *POMC^-/-^*and ADX models indicate that they are extremely similar (Figure 4H), including the PSC marker gene *Krt18* (Figure 4E and F). In summary, the loss of adrenal function whether through knockout of the *POMC* gene or through ADX leads to mobilization of PSCs and establishment of a population of pre-corticotropes followed by expansion of the mature corticotrope population. The pre-corticotropes differ from corticotropes by expression of PSC markers like *Krt18* (Figure 2H, I), and they demarcate from proliferating precursors by higher (detectable in IHF) expression of Tpit (Figure 1F, H). The pre-corticotropes are also unique with a signature of high BDNF expression together with expression of *Ntrk2* receptor and of the BDNF-dependent *Vgf* gene, suggesting an autocrine BDNF pathway loop operating in pre-corticotropes.

### BDNF is crucial for postnatal pituitary cell expansion

While the BDNF autocrine loop operating in pre-corticotropes may be important for maintenance of corticotrope lineage cells, BDNF signaling towards PSCs may be important for mobilization of stem cells and their entry into the differentiation pathway. Our previous analyses of the PSC subsets indicate that ADX leads to general mobilization of PSCs with secondary maintenance of the relevant cell types (Rizzoti, Chakravarty et al. 2023). We analyzed the *BDNF^-/-^* mice to assess whether this signalling pathway plays a role in normal pituitary development. A striking reduction of pituitary size is observed in P14 *BDNF^-/-^*mice (Figure 5C compared to 5A) but not in P10 *BDNF^-/-^*pituitary (Figure S5). A slight reduction of pituitary size is also observed for *BDNF^+/-^*pituitaries (Figure 5B) consistent with previous reports (Gururajan, Hill et al. 2015). The timing of the size difference thus appears to coincide with the postnatal phase of pituitary growth (Laporte, Vennekens et al. 2020, Sheridan, Chakravarty et al. 2025). Assessment of Ki67 labelling in *BDNF^-/-^* pituitaries shows a significant decrease of proliferating cells in the *BDNF^-/-^*pituitary consistent with a crucial role of BDNF for mobilization of postnatal PSCs and the establishment of the adult pituitary size (Figure 5D-G). This decrease in cell proliferation does not significantly affect the distribution of different pituitary lineages, with one exception, consistent with an effect that is primarily exerted by BDNF on mobilization of PSCs. Indeed, the relative abundance of somatotropes (GH, Figure 5H), lactotropes (PRL, Figure 5I), SOX2-positive PSCs (Figure 5J) and corticotropes (Tpit, Figure 5K, L) is similar in *BDNF^-/-^* compared to *BDNF^+/+^*mice, but there are less gonadotropes, particularly in the dorsal anterior lobe (LH, Figure 5M, N), consistent with previous work showing a particular dependence of this lineage on PSCs (Sheridan, Chakravarty et al. 2025).

**Figure 5.**
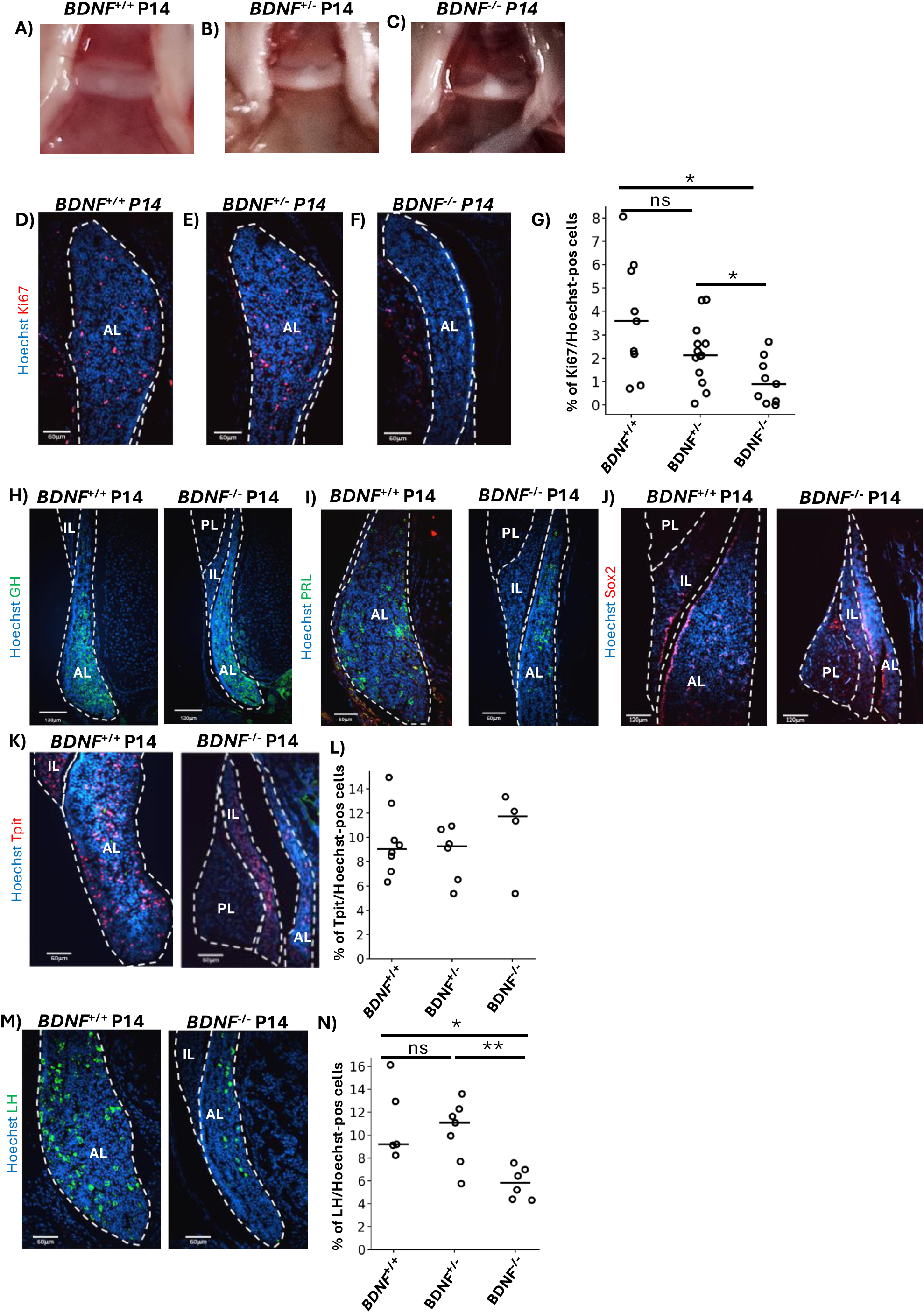
BDNF is required for postnatal expansion of the pituitary gland. (**A-C**) The *BDNF^-/-^* mice fail to expand the anterior pituitary during postnatal development (**C**) compared to *BDNF^+/+^* (**A**) mice at postnatal day 14 (P14). The *BDNF^+/-^* mice (**B**) have slightly reduced anterior pituitary. (**D-G**) The proliferative index measured by Ki67 labelling is reduced in *BDNF^+/-^* **(F)** and *BDNF^-/-^* **(E)** pituitaries compared to *BDNF^+/+^***(D)** mice. (**G**) Quantitation of the number of Ki67 positive cells in pituitaries of the indicated genotypes (*BDNF^+/+^* n=9 from 3 mice, *BDNF^+/-^* n=13 from 4 mice & *BDNF^-/-^* n=9 from 3 mice). (**H-K**) Cell differentiation is not severely impaired by *BDNF* gene knockout as revealed by immunostaining for GH (**H**), prolactin (PRL) (**I**), Sox2 (**J**) and Tpit (**K, L**). (**L**) Quantitation of the number of Tpit positive cells in pituitaries of the indicated genotypes (*BDNF^+/+^* n=8 from 2 mice, *BDNF^+/-^*n=6 from 3 mice & *BDNF^-/-^* n=4 from 2 mice). (**M**) Quantitation of the number of LH positive cells in pituitaries of the indicated genotypes (*BDNF^+/+^* n=5 from 3 mice, *BDNF^+/-^* n=7 from 4 mice & *BDNF^-/-^* n=6 from 3 mice). (**M, N**) The only lineage affected by *BDNF* knockout is the gonadotropes marked by LH expression. Statistical significance was measured by paired two-sided T-test: * p≤0.05, **p≤0.01.

In summary, BDNF is a major signal for expansion of the pituitary gland during the postnatal period (Figure 6). A likely source of BDNF during the postnatal phase of pituitary development may well be the somatotropes and lactotropes that express *BDNF* (Figure 3C) and that expand independently of PSCs during the same period (Sheridan, Chakravarty et al. 2025).

**Figure 6.**
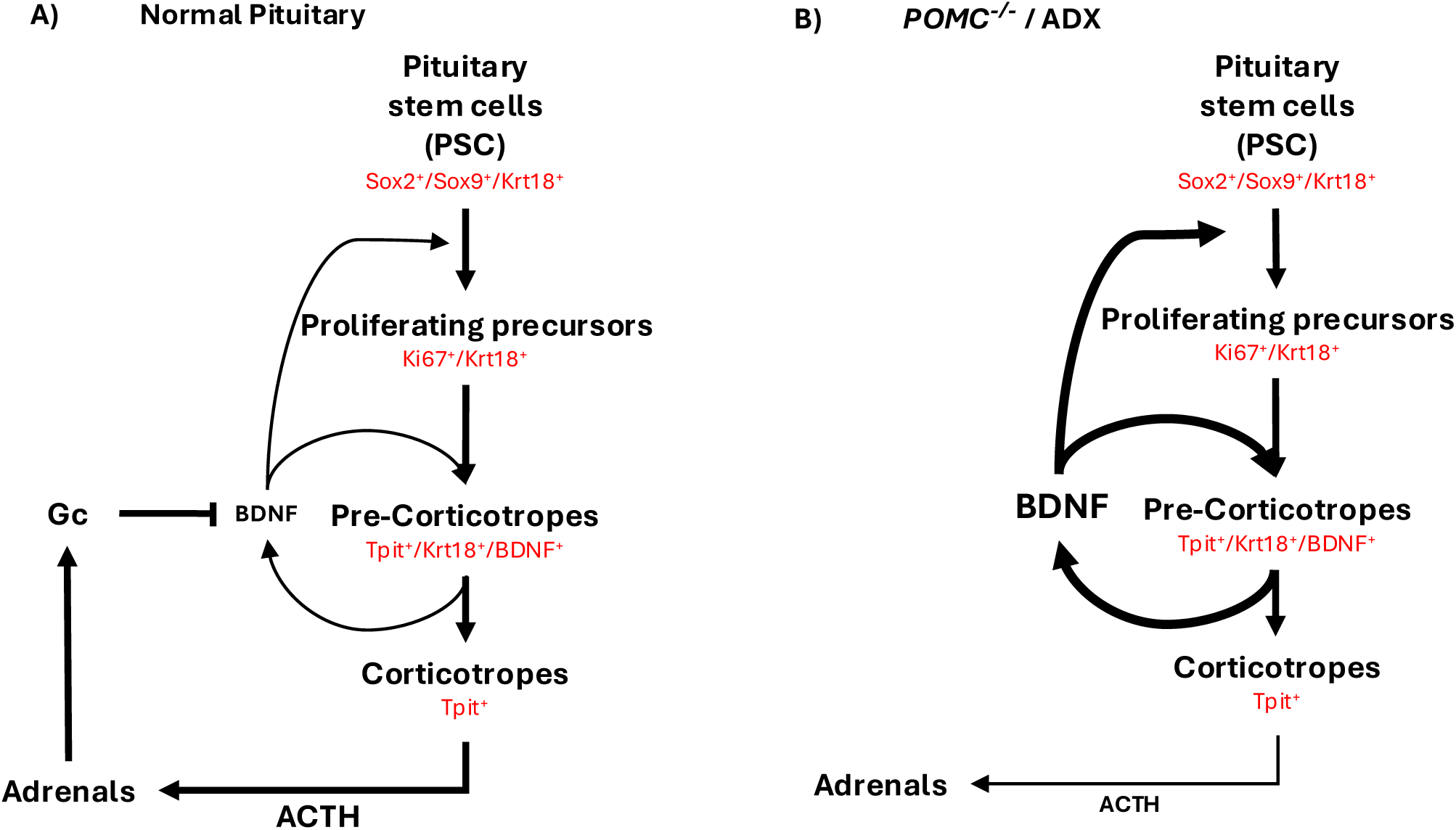
Model of BDNF signalling in control of pituitary stem cell proliferation and differentiation in normal (A), and *POMC^-/-^* and ADX (B) pituitaries. (**A**) In normal pituitary development, BDNF is an important signal for postnatal entry of pituitary stem cells (PSCs) into proliferation and differentiation. It is presumably important for pre-corticotrope maintenance, but these cells do not accumulate in the normal pituitary. Pre-corticotrope BDNF expression is repressed by glucocorticoids (Gc). (**B**) In *POMC^-/-^* and ADX mice, pre-corticotrope BDNF expression is de-repressed because of deficient Gc production from the adrenals and these cells accumulate in significant number. Pre-corticotrope-derived BDNF also stimulates entry of PSCs into proliferation and differentiation.

## Discussion

The present work identified and defined the properties of a novel pituitary cell type, the pre-corticotropes. Although their existence was a priori likely, their crucial role in pituitary cell homeostasis and their origin was not. Their massive expansion in the *POMC^-/-^* model and their dependence on lineage specification by Tpit together with their sensitivity to Gc regulation enabled the discovery of a critical signalling pathway for homeostasis of the pituitary-adrenal axis. Pre-corticotropes do not accumulate in the normal pituitary and indeed there are so few that they can only be recognized *a posteriori*. Since their maintenance is repressed by Gc’s, they arise and expand in mice that are deficient in Gc production either caused by *POMC* gene inactivation (*POMC^-/-^*) or by ADX. Further, their establishment is lineage-dependent, and more specifically dependent on the lineage specifying transcription factor Tpit. Pseudotime analyses of scRNA-seq data together with lineage tracing in the Sox9iresGFP model indicate that pre-corticotropes originate from PSCs that are induced to proliferate. Proliferating PSCs appear as transit amplifying cells that retain some PSC markers while they gain corticotrope markers. These Ki67-positive cells do not accumulate significantly in both *POMC^-/-^* and ADX models. The reason why pre-corticotropes accumulate more in aged *POMC^-/-^* compared to ADX model is not clear.

The analysis of ligand-receptor pairs expressed in pre-corticotropes compared to mature corticotropes identified the BDNF pathway as a candidate for the pituitary cell response to end-organ ablation. Failure of *BDNF^-/-^* mice to undergo pituitary cell expansion in the postnatal period clearly supports the model that BDNF is the crucial signal to engage postnatal PSCs into proliferation and differentiation. This activity does not seem to be lineage specific but rather reflect a direct effect of BDNF on PSCs that express the BDNF receptor *Ntrk2*. While BDNF is expressed at low levels in multiple differentiated pituitary cells, its high expression in pre-corticotropes together with Dex-dependent repression of this expression supports the model that in *POMC^-/-^* pituitaries, it is pre-corticotrope-derived BDNF that stimulates PSCs into proliferation followed by BDNF-dependent maintenance of *Ntrk2*-expressing pre-corticotropes. Maintenance of pre-corticotropes thus appears to involve a BDNF autocrine loop. Collectively, the data support a crucial role of pre-corticotropes in maintaining the balance between pituitary corticotropes and adrenal function (Figure 6). In contrast to the effect of BDNF on proliferation of PSCs, the homeostatic function of pre-corticotropes within the corticotrope axis is ensured by the dependence of this cell type on its lineage specification by Tpit. It is thus noteworthy that there is not only failure to expand the pre-corticotrope cell population but also failure to engage PSCs into proliferation in the *POMC^-/-^;Tpit^-/-^* and *POMC^-/-^;Tpit^+/-^*pituitary.

In summary, the present work suggests that the BDNF pathway has a unique role in establishment of the adult population of the pituitary gland during the postnatal period when gland function is adapted to organismal growth and that the same signalling pathway is crucial for adaptation in conditions of major stress on the HPA axis such as exerted by ADX. In contrast, we have no indication of a similar role for the BDNF pathway in the initial formation of the pituitary gland during the foetal period: we must therefore conclude that other mechanisms are involved in early expansion and differentiation of the pituitary primordium, Rathke’s pouch. The Wnt pathway was involved in these processes (Potok, Cha et al. 2008, Russell, Lim et al. 2021).

## Acknowledgments

We are grateful to Ludovic Malet for assistance in analysis of scRNA-seq data, to Sarah Boissel for next-generation sequencing, to Sophie Wood from the Francis Crick surgical service platform for performing adrenalectomies, to Anabelle Bouchard for the histological sample preparation and to Valerie Magoon for expert secretarial assistance. This work was supported by grants to JD from the Canadian Institutes of Health Research (FDN-154297) and the Digital Research Alliance of Canada (zmv-495 553), to MER from National Institute of Neurological Disorders and Stroke of the National Institutes of Health (R01 NS135884), and to RLB from Cancer Research UK (FC001107), the UK Medical Research Council (FC001107), and the Wellcome Trust (FC001107).

## Author Contributions

Conceptualization: Kevin Sochodolsky, Konstantin Khetchoumian, Aurelio Balsalobre, Karine Rizotti, Jacques Drouin

Methodology: Kevin Sochodolsky, Karine Rizotti, Aurelio Balsalobre, Jacques Drouin

Investigation: Kevin Sochodolsky, Ryan M. Feeley, Karine Rizotti, Probir Chakravarty, Aurelio Balsalobre, Jacques Drouin

Visualization: Kevin Sochodolsky, Karine Rizotti, Jacques Drouin

Funding acquisition: Margaret E. Rice, Robin Lovell-Badge, Jacques Drouin

Project administration: Jacques Drouin

Supervision: Karine Rizotti, Aurelio Balsalobre, Jacques Drouin

Writing – original draft: Jacques Drouin

Writing – review & editing: Kevin Sochodolsky, Karine Rizotti, Jacques Drouin

## Declaration of interests

Authors declare that they have no competing interests.

## STAR Methods

### Resource availability

Lead contact: Jacques Drouin

#### Materials availability

Data and code availability: The datasets analyzed in this study are available from the Gene Expression Omnibus (GEO) repository under the following accession numbers: GSE316726, GSE217648, GSE313979.

### Experimental Model and Subject Details

#### Mice

The *POMC^-/-^* mice originally generated by Yaswen et al. (Yaswen, Diehl et al. 1999) were obtained from Jackson Labs (B6.129X1-Pomctm2Ute/J strain, stock number 008115) and the *Tpit^-/-^*mice were described previously (Pulichino, Vallette-Kasic et al. 2003, Pulichino, Vallette-Kasic et al. 2003). Animal experimentations were approved by the IRCM Animal Care and Use Committee in conformity with regulations of the Canadian Council on Animal Care.

All animal experiments carried out in London (Figure 4) were approved under the UK Animals (Scientific Procedures) Act 1986 under the project license PP8826065 and by the Francis Crick Animal Welfare and Ethical Review Body (AWERB). Seven-week-old C57BL/6Jax mice were subjected to bilateral adrenalectomies (ADX), and pituitaries harvested 4 days after surgery, as described (Rizzoti, Chakravarty et al. 2023).

All animal handling procedures for *BDNF^+/-^* and *BDNF^-/-^*mice and their wildtype littermates at the NYU Grossman School of Medicine were conducted in accordance with the National Institutes of Health guidelines and approved by the New York University Grossman School of Medicine Animal Care and Use Committee. Founders for this colony were provided by Prof. Francis S. Lee at Weill Cornell Medicine. Founders were *BDNF ^+/-^* females (B6.129S4-Bdnf^tm1Jae^/J) (Ernfors, Lee et al. 1994) crossed with males (Stock number 000664) from Jackson Labs, and *BDNF^-/-^* mice were generated by crossing *BDNF^+/-^*mice from this colony. This breeding paradigm successfully produced BDNF knockouts that lived to experimental age and age matched wildtypes; however, they were produced rarely. All mice were genotyped by the Genotyping Core Laboratory at NYU Langone Health at ≤ P7 using published protocol (Jackson Labs, protocol 22247).

### Method Details

#### Pituitary dissection and preparation

Pituitaries from available *BDNF^-/-^* mice, P10-P14, were isolated, along with those of WT and BDNF^+/-^ littermates of the same age. Before dissection, mice were deeply anesthetized using isoflurane. After the brain was removed, the skull containing the pituitary was transferred to 4% paraformaldehyde (PFA) in phosphate buffered saline (pH 7.3) for overnight fixation, then stored in 70% ethanol until further processing, as described (Bilodeau, Roussel-Gervais et al. 2009).

#### Immunofluorescence (IHF) staining

Processing of PFA-fixed, paraffin-embedded pituitaries and immunohistofluorescence were described previously (Bilodeau, Roussel-Gervais et al. 2009).

#### RNAscope

*Pomc* (# 314081) and *Vgf* (# 517421) mRNAs were visualized on cryosections of PFA-fixed pituitaries using RNAscope, following the manufacturer’s protocol.

#### scRNA-seq

Mouse pituitary cells were dispersed and used in scRNA-seq as described (Khetchoumian, Sochodolsky et al. 2024). ADX mice pituitary cells were dispersed and used for scRNA-seq as described (Rizzoti, Chakravarty et al. 2023).

The *POMC^+/+^* and *POMC^-/-^* scRNA-seq data analyses, filtering, normalization and scaling, dimensionality reduction and standard unsupervised clustering were performed using the Seurat R package v3.1 (Satija, Farrell et al. 2015). Low quality cells were excluded using these parameters: 1500 > nFeature counts > 4000; nCount RNA > 10,000; 5 > % mitochondrial DNA > 0.5. The samples were then combined using the merge function in Seurat. Transcript counts were normalized for total read depth. Cell clusters were identified by principal component analysis (PCA) and visualized by uniform manifold approximation and projection (UMAP) embedding (McInnes, Healy et al. 2018) and unsupervised graph-based clustering with the Louvain algorithm (Blondel, Guillaume et al. 2008, Lancichinetti and Fortunato 2009) with a resolution of 0.8. UMAP identified cell clusters were annotated based on differential expression of key pituitary marker genes as previously described (Khetchoumian, Sochodolsky et al. 2024).

Single RNAseq datasets from SOX9iresGFP sorted cells in ADX and sham operated mice 48h00 after surgeries have been described previously (Rizzoti, Chakravarty et al. 2023). Here we generated equivalent datasets from dissociated whole pituitaries harvested from 3 x 2 7-week-old C57Bl6 males that had undergone either bilateral adrenalectomies or sham-surgery. Raw Reads were aligned to the mm10 transcriptome using CellRanger (version 3.0.2) and STAR (version 2.5.1) and reported cell-specific gene expression count matrices. All subsequent analysis were performed using the Seurat (version 4) package in R (version 4.1). Each sample was subjected the following QC thresholds: Genes were considered "expressed" if the estimated log10 count was at least 0.1; Cells were removed if cells expressed less than 50 genes; Cells for which mitochondrial genes made up greater than 3 standard deviations from the mean of the mitochondrial expressed genes per sample. Doublets were identified using DoubletFinder (version 2.0.4) and scDblFinder (version1.20.2); cells called Doublets by both methods were removed.

PCA decomposition was performed per sample using the first 20 components to construct uniform manifold approximation and projection (UMAP) plots per samples.

Whole and sorted samples were integrated using Seurat’s rpca integration method to consider the differences in cellular composition between whole and sorted samples, using the top 3000 variable genes and the first 14 principal components as determined using intrinsicDimensions (version 1.2 package) to construct the UMAP.

#### Pseudotime trajectory analysis

Single-cell transcriptomic trajectory was inferred using Monocle 3 (v1.4) implemented in R (v4.4.3) (Cao, Spielmann et al. 2019). Preprocessed single-cell RNA-seq count matrices and associated cell metadata from clusters for pituitary stem cells, proliferating cells and the Tpit expressing cells from the *POMC^-/-^*sample were imported into Monocle as a *cell_data_set* object. Dimensionality reduction was performed using PCA followed by UMAP as implemented in Monocle 3. A k-nearest neighbor graph was constructed in the reduced-dimensional space using the *learn_graph()* function, enabling reconstruction of cell-state transitions. To infer pseudotime ordering, a biologically informed root node was selected based on known marker gene expression corresponding to the earliest cellular state. Pseudotime values were then assigned to each cell using the *order_cells()* function.

#### Ligand-receptor inference

Ligand-receptor inference was performed using CellChat v2 (Jin, Plikus et al. 2025) package for R (v4.4.3). Analysis was performed for every cell cluster from the Seurat analysis with more than 10 cells. *POMC^+/+^* and *POMC^-/-^* samples were analysed together and Sham and ADX were analysed together. The selected ligand-receptor interacting pairs were visualised using Dotplot representation generated by the Seaborn statistical data visualization tool (https://seaborn.pydata.org/) in Python (v3.8). Dot size inversely correlates with the p-value and colour indicates the interaction probability score.

#### Violin plots and gene expression UMAPs

Violin plots and gene expression UMAPs from scRNA-seq data were generated with Loupe Browser 8.0.0 from 10X Genomics (https://www.10xgenomics.com/support/software/loupe-browser/latest) using the LogNorm scale value.

#### Differential gene expression analysis

Differential gene expression analysis was performed using Loupe Browser 8.0.0 from 10X Genomics with the LogNorm scale. The cutoff for the differentially expressed genes is a log2(fold change) above 1 or below -1 with a p-value smaller than or equal to 0.05. The data was then visualised using the *plt.scatter()* function of the Matplotlib library (https://matplotlib.org/) in Python (v3.8). Heatmaps were generated with Seaborn statistical data visualization tool (https://seaborn.pydata.org/) in Python (v3.8).

### Quantification and statistical analysis

Details of statistical analyses are provided in the figure legends.

### Key resources table

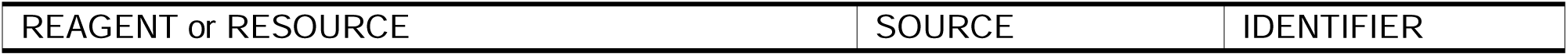
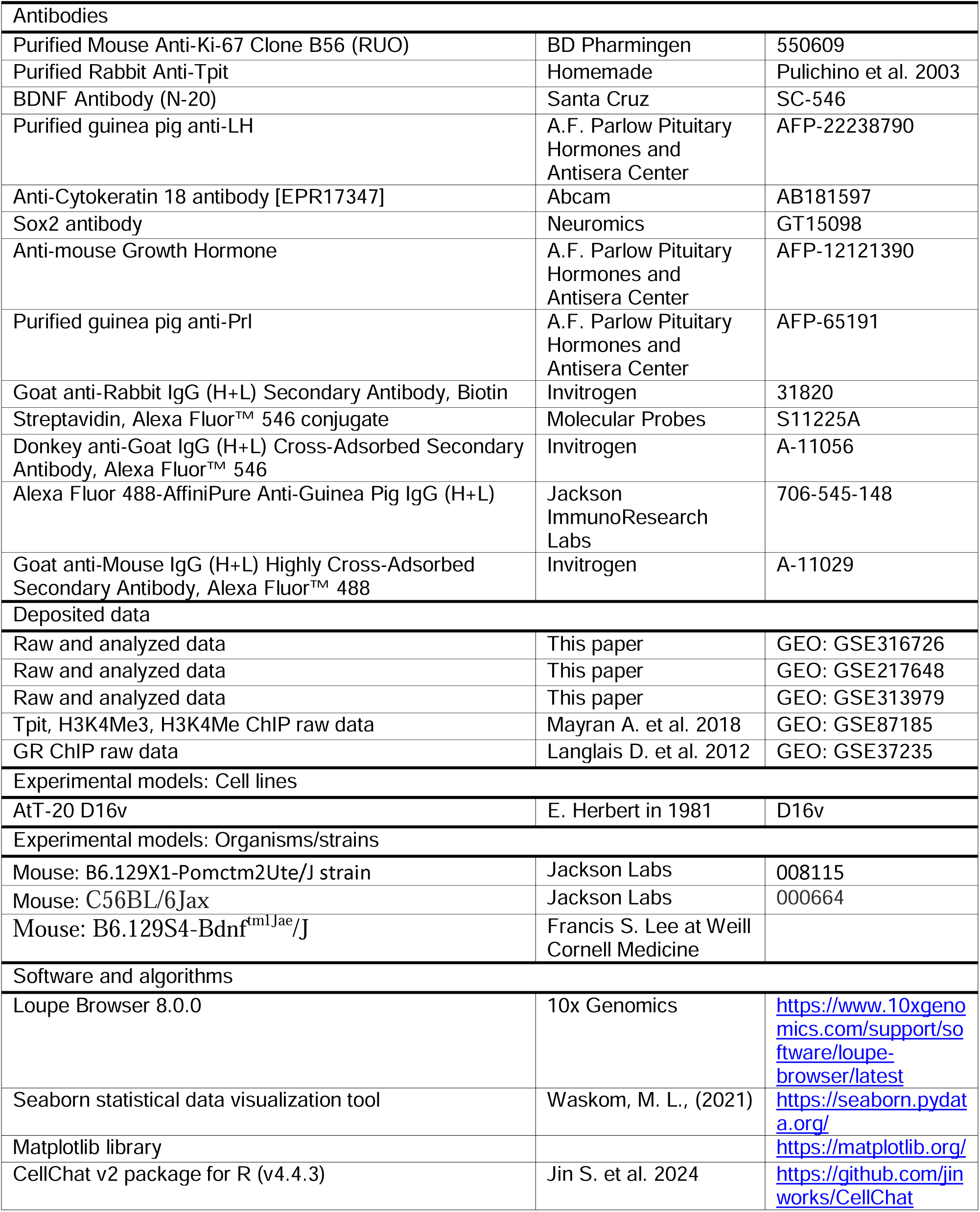
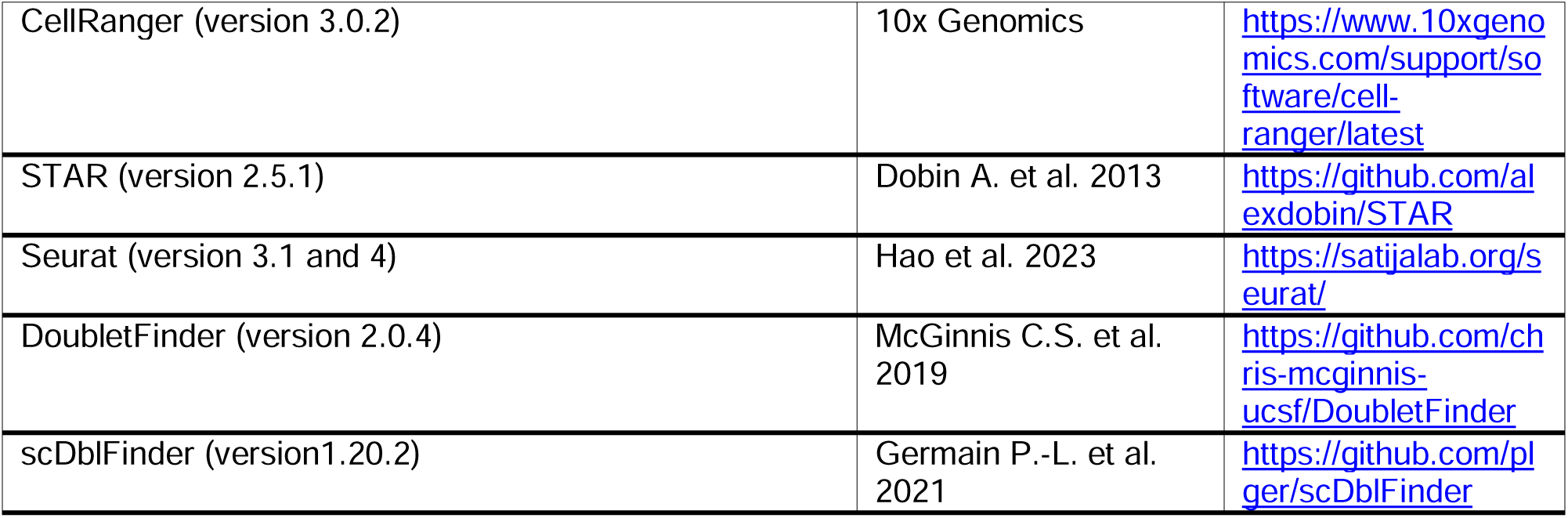

## Supplemental Information

Document S1. Figures S1-S5

Table S1. Excel file of differentially expressed genes from POMC^-/-^ pre-corticotropes compared to POMC^+/+^ corticotropes related to Figure 3A.

## Notes

### Competing Interest Statement

The authors have declared no competing interest.

