## Supplementary Figures for "BDNF Regulates Pituitary Stem Cell Engagement towards precursor state"

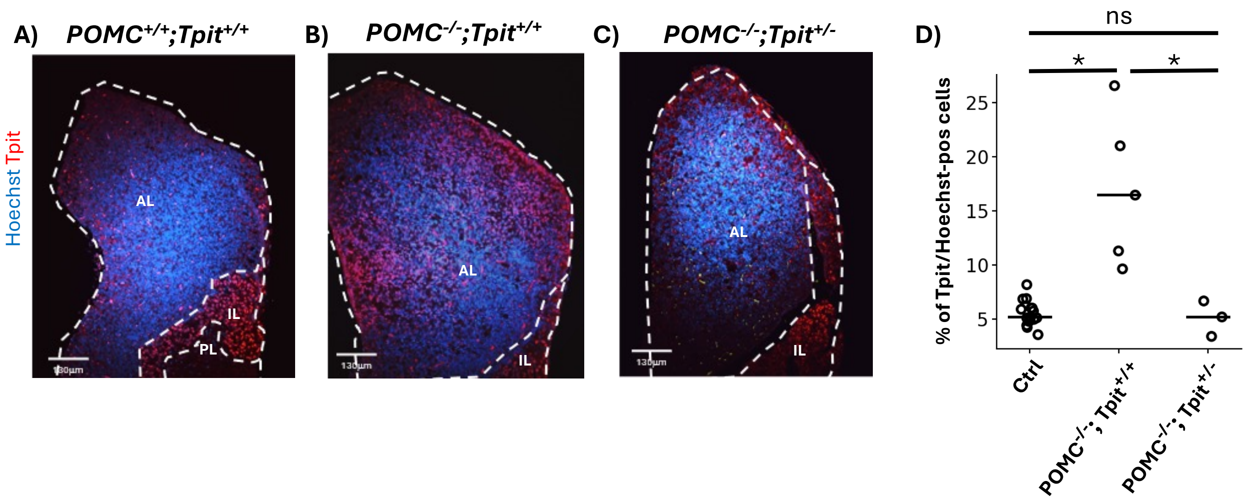


**Figure S1** *Tpit* haploinsufficiency prevents corticotrope hyperplasia. (**A-C**) Tpit staining of pituitary sections from mice of the indicated genotypes show that Tpit-positive corticotrope hyperplasia present in *POMC*$^{-/-}$*;Tpit*$^{+/+}$ pituitary (**B**) compared to control (**A**) is prevented by *Tpit* gene haploinsufficiency (**C**). (**D**) Quantitation of Tpit-positive cells in pituitary sections of the indicated genotypes (Ctrl n=15 from 6 mice, *POMC*$^{-/-}$*;Tpit*$^{+/+}$ n=5 from 4 mice & *POMC*$^{-/-}$*;Tpit*$^{+/-}$ n=3 from 3 mice). Statistical significance as measured by paired T-test: * $p \leq 0.05$.

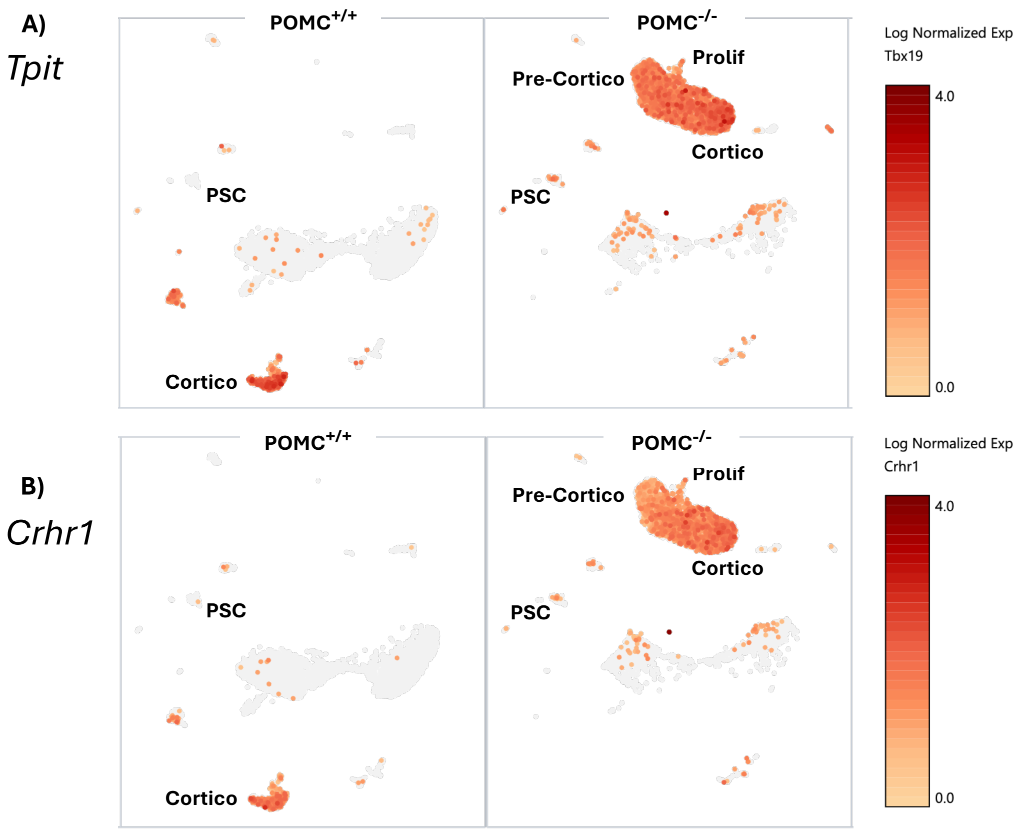


**Figure S2** (**A**) UMAP representation of *Tpit* gene expression in scRNA-Seq analyses of the pituitary from *POMC*$^{-/-}$ mice. (**B**) UMAP representation of CRH receptor 1 (*Crhr1*) gene expression in scRNA-Seq analyses of the pituitary from *POMC*$^{-/-}$ mice.

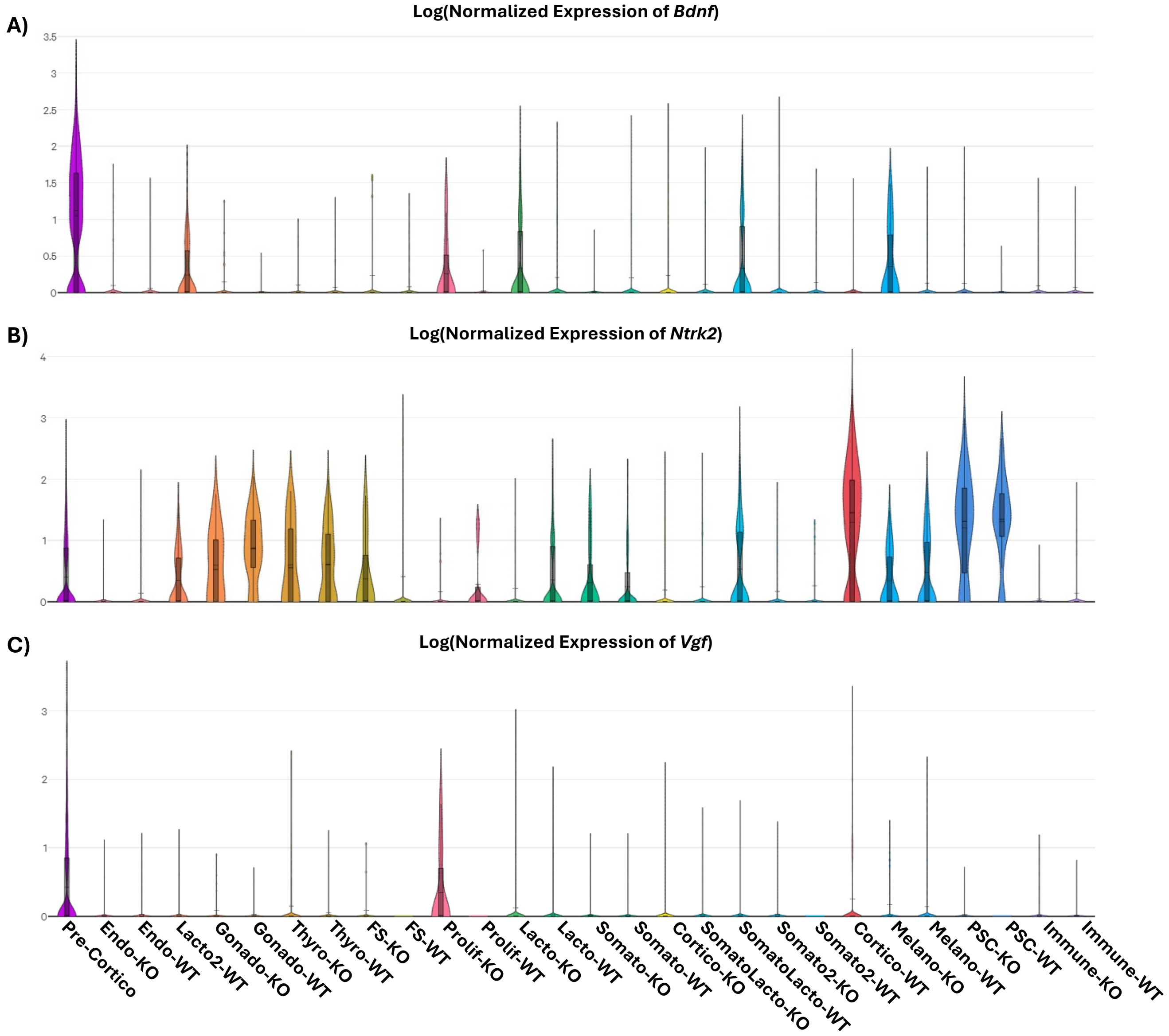


**Figure S3** Relative expression of BDNF pathway genes in different cell clusters from pituitary scRNA-Seq analyses. (**A**) Violin plots showing relative expression of *BDNF* gene in cell clusters from pituitary scRNA-Seq analyses of control and *POMC*$^{-/-}$ pituitaries. (**B**) Relative expression of *Ntrk2* gene in different clusters from pituitary scRNA-Seq analyses. (**C**) Violin plots showing relative expression of *Vgf* gene in cell clusters from scRNA-Seq analyses of control and *POMC*$^{-/-}$ pituitaries.

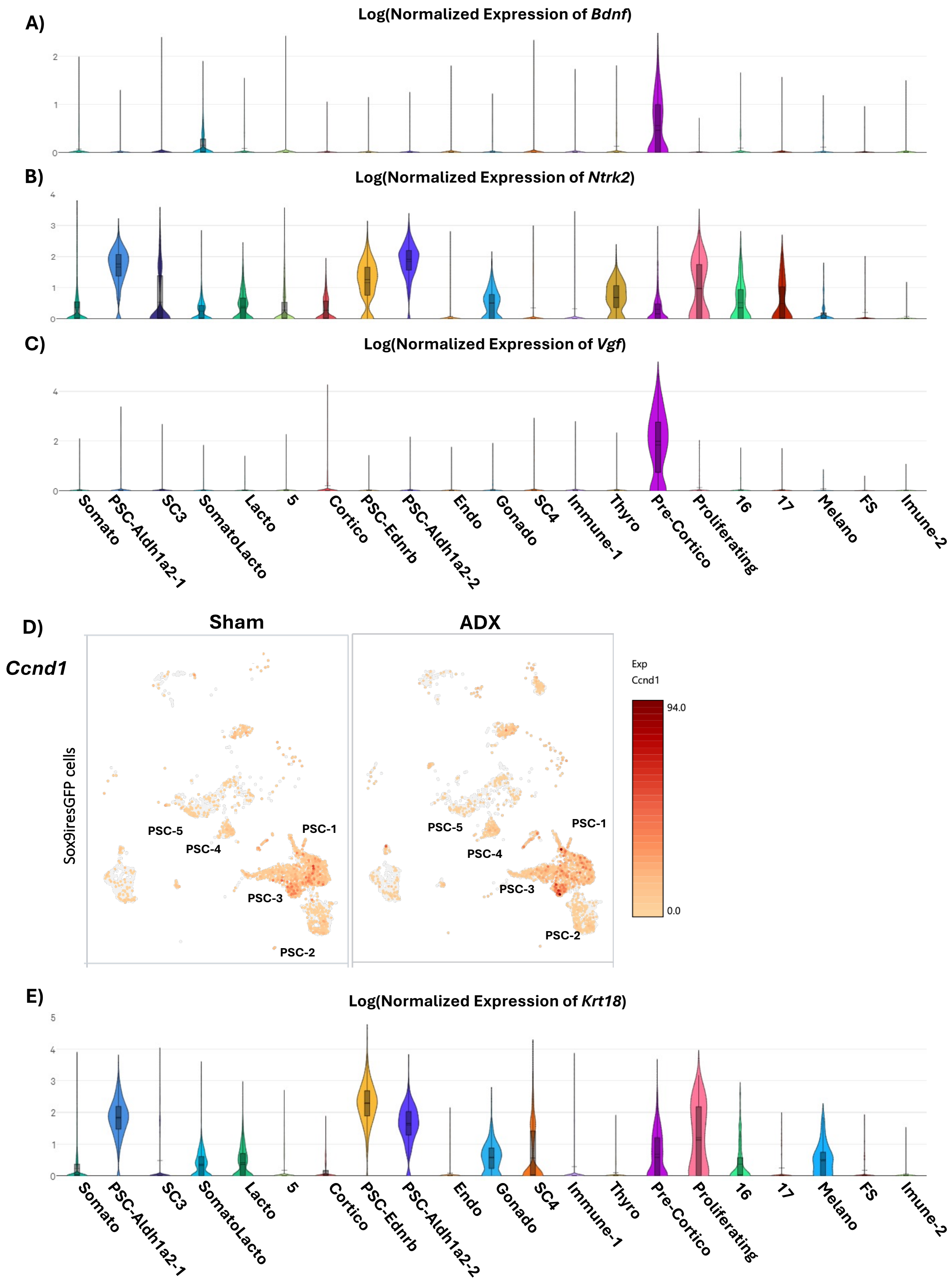


**Figure S4** Relative expression of BDNF pathway genes in ADX compared to SHAM operated mice. (**A**) Relative expression of *BDNF* gene in different cell clusters from pituitary scRNA-Seq analyses of Sham and ADX mice. (**B**) Relative expression of *Ntrk2* gene in different cell clusters from pituitary scRNA-Seq analyses of Sham and ADX mice. (**C**) Violin plots showing relative expression of *Vgf* gene in cell clusters from scRNA-Seq analyses of Sham and ADX mice. (**D**) UMAP representation of Cyclin D1 (*Ccnd1*) gene expression in scRNA-Seq analyses of pituitary Sox9-positive cells from sorted Sox9iresGFP mice. (**E**) Relative expression of *Krt18* gene in different cell clusters from pituitary scRNA-Seq analyses of Sham and ADX mice.

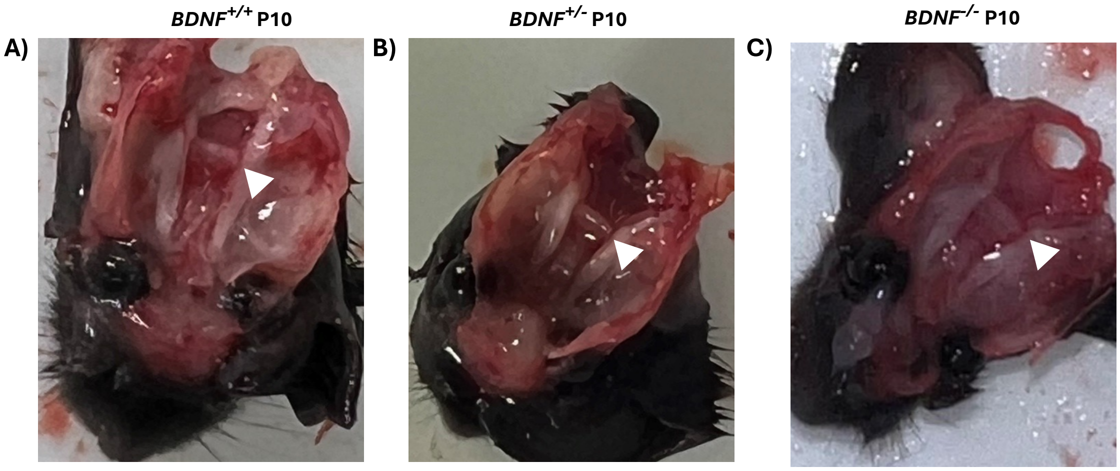


**Figure S5** Photographs of in situ pituitaries for mice of the indicated genotypes at postnatal day 10 of development (P10).
